# Modeling How Community Assembly Alters the Functioning of Ecosystems

**DOI:** 10.1101/2020.02.10.942656

**Authors:** Thomas Koffel, Colin T Kremer, K. Bannar-Martin, S.K. Morgan Ernest, Nico Eisenhauer, Christiane Roscher, Juliano Sarmento Cabral, Mathew A. Leibold

**Affiliations:** W.K. Kellogg Biological Station, Michigan State University, 3700 E Gull Lake Dr., Hickory Corners, MI 49060. U.S.A.; W.K. Kellogg Biological Station, Michigan State University, 3700 E Gull Lake Dr., Hickory Corners, MI 49060. U.S.A; Department of Ecology and Evolutionary Biology, University of California Los Angeles, 612 Charle E. Young Drive East, PO Box 957246, Los Angeles, CA 90095-7246 USA.; Department of Wildlife Ecology and Conservation, University of Florida, Gainesville, FL 32611, USA; German Centre for Integrative Biodiversity Research (iDiv) Halle-Jena-Leipzig, Deutscher Platz 5e, 04103 Leipzig, Germany; Institute of Biology, Leipzig University, Deutscher Platz 5e, 04103 Leipzig, Germany.; Helmholtz Centre for Environmental Research (UFZ), Department Physiological Diversity, Permoserstrasse 15, 04318 Leipzig, Germany; German Centre for Integrative Biodiversity Research (iDiv) Halle-Jena-Leipzig, Deutscher Platz 5e, 04103 Leipzig, Germany.; Ecosystem Modeling, Center for Computational and Theoretical Biology (CCTB), University of Würzburg, Emil-Fischer-Str. 32, 97074 Würzburg, Germany.; Department of Biology, University of Florida, Gainesville, FL 32611, USA.

## Abstract

Although the effects of species richness on ecosystem functioning have been extensively studied, there is increased interest in understanding how community assembly in general might alter the functioning of ecosystems. We focus on two complementary approaches for evaluating how community assembly influences ecosystem function (here, productivity). The first quantifies the relative importance of complementarity and selection by contrasting monocultures with polycultures. The second identifies the effects of species losses and/or gains relative to the baseline polyculture, as well as the indirect effects on other species’ productivity. We evaluate and contrast these two approaches, using simulated communities structured by different, known competition mechanisms, where species compete for different resources and experience varying levels of environmental heterogeneity. We find that the metrics provided by these approaches can jointly discriminate the mechanisms of competition driving productivity. We then apply our methods to data from a long-term biodiversity-ecosystem experiment (the Jena Experiment) and find that the data do not correspond to any of the competition scenarios we modeled. We address two additional possible complications: facilitation by nitrogen fixing plants, and non-equilibrium behavior during community assembly, and find that a combination of resource competition and facilitation by nitrogen fixing plants is the more likely explanation for the results obtained at Jena.

## Introduction

Community assembly drives shifts in community structure (i.e., species richness and composition) which can have important consequences for the functioning of ecosystems. While the impact of changes in species richness on ecosystem function (commonly called biodiversity-ecosystem function, or BEF, effects; reviewed by Cardinale et al. 2012, Hooper et al. 2012, Tilman et al. 2012) are the best-studied, the importance of all of community assembly’s impacts on community structure (referred to here as community assembly-functioning of ecosystem, or CAFE, effects) is increasingly recognized (e.g. Ptacnik et al. 2010, Thompson and Gonzalez 2016, Hodapp et al. 2016, Leibold et al. 2017, Bannar-Martin et al. 2018). Drivers of community assembly include species interactions, disturbance, dispersal, environmental heterogeneity, and landscape effects that can lead to changes in community composition, without necessarily affecting species richness (Brown et al. 2001). Community assembly is thus complex and depends on multiple factors and processes that may also consequently affect ecosystem function (e.g. Thompson and Gonzalez 2016, Hodapp et al. 2016, Leibold et al. 2017). Therefore, it is community assembly processes as a whole that ultimately link community composition, including species richness, to ecosystem attributes.

How can we study these community assembly effects on ecosystem function? One approach would be to comprehensively study the detailed interactions among the species involved and theoretically analyze these interactions to fully disentangle how community assembly affects ecosystem attributes. This approach, however, is logistically demanding and likely to give answers that are system specific. An alternative is to develop metrics that parse the effects of various components of community assembly on ecosystem attributes. Useful measures should be general enough to allow cross-system comparisons and simple enough that they reduce the full complexity of communities, while remaining sufficiently connected to real community features to be informative. In this paper we focus on two of several methods that provide complementary information, the Complementarity-Selection method and the Price equation components (e.g. Kirwan et al. 2009, Bell et al. 2009, Hector et al. 2009, Bannar-Martin et al. 2018).

The Complementarity-Selection method (originally called the ‘additive partitioning method’) offers insights into the ecosystem function of polycultures (communities) by comparing them with null models of hypothetical polycultures where species’ functions are proportional to their function in monoculture (Loreau and Hector 2001). Total ecosystem functioning is partitioned into the function arising from: (i) selection effects (which are positive when polycultures are dominated by species that are high functioning in monoculture), and (ii) complementarity effects (which are positive when species contribute more function to polycultures than expected given their functioning in monoculture, due to interactions like niche partitioning or facilitation). Both of these effects can also be negative, if low-functioning species dominate or species interactions reduce function. It is generally thought that selection effects occur due to environmental filtering, which selects for more productive species that are better suited to the local environment. Complementarity is typically related to niche partitioning that results in enhanced coexistence and increased resource-use efficiency. This approach is well-suited to the design of most BEF experiments, as they quantify the function (productivity) of monocultures of species used to form higher-diversity polycultures. However, this approach does not allow direct comparison of communities of varying composition and structure.

Another approach for understanding how the ecosystem function of communities (polycultures) relates to community assembly, the Price equation method, focuses directly on comparing pairs of communities (Fox 2006). Here, differences in total function between communities are related to changes in composition and individual species’ functions, as quantified by calculating ‘Price components’ (Fox 2006, Fox and Kerr 2012, Bannar-Martin et al. 2018). Price components can be used flexibly to make different types of inferences depending on choices about (i) which communities are compared, and (ii) which of several arrangements of Price components are examined (Bannar-Martin et al. 2018). For this paper, we focus on (i) comparing communities with the most diverse species pool to communities with lower levels of richness, and (ii) using a three-part Price component arrangement that separates the functional effects of species gains (‘SG’), species losses (‘SL’) and the differences in density/function among species present in both communities being compared (the Context Dependent Effect or ‘CDE’). In Bannar-Martin et al. (2018) these are called the Community Assembly components; we use this terminology in the rest of this paper. Price equation approaches like the Community Assembly analysis can be applied more generally than the Complementarity-Selection method, and used to examine how <u>diverse</u> changes in community structure (including species richness, taxonomic turnover, and species’ abundances, function, or traits; Fox & Harpole 2008) alter ecosystem functioning. However, their use is restricted to ecosystem functions that can be calculated as the sum of contributions of their constituent species (Fox 2006).

We study how these two approaches might be combined, and if jointly they can enhance our understanding of the mechanisms underlying the functioning of communities. Our ultimate goal is to better understand real-world communities. However, we begin by applying these methods to predictions from a community model we constructed based on resource-competition theory (Tilman 1982, Tilman et al. 1997, Schreiber and Tobiason 2003, Gross and Cardinale 2007, Hodapp et al. 2016, Koffel et al. 2016) that can emulate typical BEF experiments. We applied the model to various assembly scenarios to consider a wide range of possible ways resource competition may occur and affect community assembly (including competition for complementary vs. antagonistic resources with homogeneous or heterogeneous resource supply at the local scale). We then apply the Complementarity-Selection approach (Loreau and Hector 2001) and the Community Assembly approach (Bannar-Martin et al. 2018) to the results of the models to assess if these methods can be used to distinguish among the various known assembly scenarios. Next, turning from theoretical results, we consider data from the long-term BEF experiment conducted in grasslands at Jena (Roscher et al. 2004, 2005; Weigelt et al. 2010) and find that the data are not consistent with any of our initial assembly scenarios. This led to a set of *a posteriori* hypotheses, which we identify and address separately (consistent with recommendations for improving transparency and reproducibility in biology; e.g. Ihle et al. 2017, Nichols et al. 2019). Specifically: in these real-world communities there may be 1) inadequate time to achieve steady-state competitive outcomes and/or 2) facilitation (e.g. due to the effects of nitrogen fixing plants). We explore these possibilities by adapting our model, and find support for the facilitation scenario. Put together, these results show how the joint use of the Complementarity-Selection and Community Assembly approaches make it possible to discriminate the mechanisms underlying the functioning of ecological communities.

## Methods

To model community assembly in simulated BEF experiments, we first considered a flexible version of the basic resource competition model for two resources as initially developed by MacArthur (1972) and further elaborated by Tilman (1982):

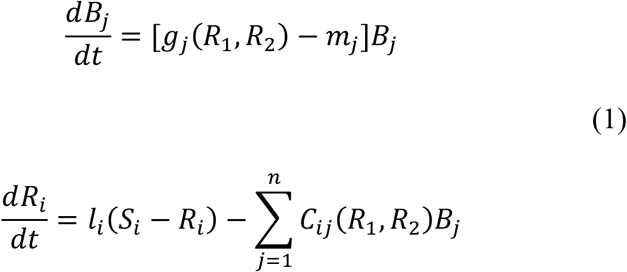

This model describes the population dynamics of *n* species *j* (whose biomasses are denoted *B_j_*) that compete for two resources, *R*_1_ and *R*_2_. Here *g_j_*(*R*_1_, *R*_2_) and *m_j_* respectively determine species *j*’s per capita growth and mortality rates, *S_i_* is the supply of resource *R_i_*, which is diluted at rate *l_i_*, and *C_ij_* (*R*_1_, *R*_2_) is the *per capita* consumptive impact of *B_j_* on *R_i_*. Species *j*’s interactions with the two resources can be divided in two parts: (1) its requirements (through the *per capita* growth rate *g_j_*) and; (2) its impacts on the resources (through the *per capita* consumptive impacts *C_ij_*).

Competitive scenarios: We considered a range of competitive scenarios, classified by the types of resources involved, including resources that are ‘strongly complementary’ (virtually ‘essential’, *sensu* Tilman 1982) vs. ‘weakly complementary’ and ‘weakly antagonistic’ (these two cases correspond closely to ‘substitutable’ in Tilman’s terminology, the perfectly substitutable model has undesirable mathematical properties for our purpose) vs. ‘strongly antagonistic’. To implement this variation in resource types we used a flexible form of the growth rate *g_j_*(*R*_1_, *R*_2_) that can generate all these scenarios by tuning a single parameter *α* (Schreiber and Tobiason, 2003; See Appendix A).

Fig. 1 illustrates how each of these resource types affects trade-off relations among minimum resource requirements for the two resources. Previous work has almost exclusively focused on the strongly complementary scenario (e.g. Tilman et al. 1997, Gross and Cardinale 2007, Hodapp et al. 2016); however, we found that different types of competition lead to different outcomes of community assembly, associated with distinctive signatures in terms of Complementarity-Selection effects and Community Assembly components (see Box 1 and ‘Results’ section).

**Figure 1.**
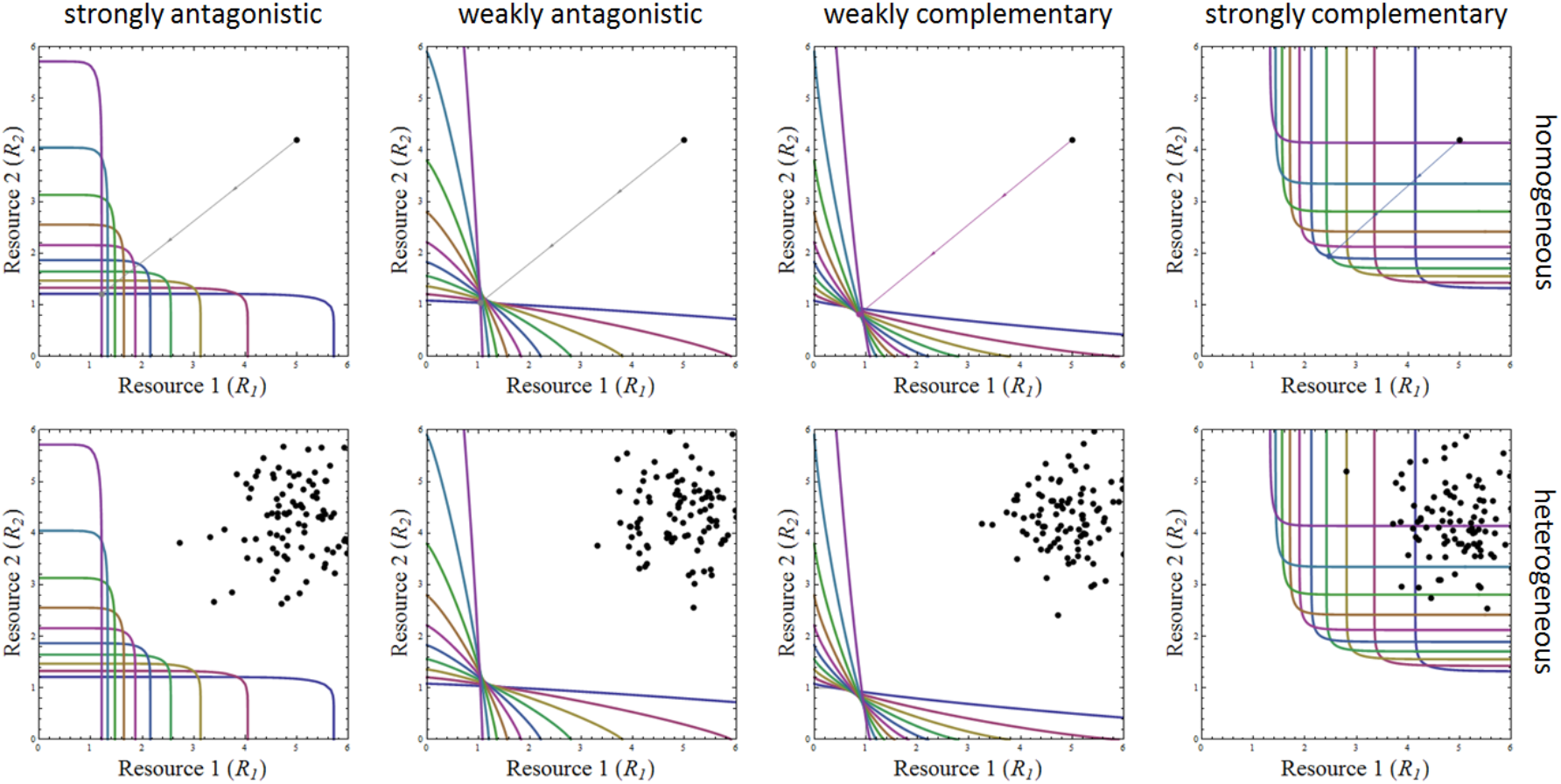
Zero Net Growth Isoclines (ZNGIs; colored, thick) and supply points (black dots) for each of the eight resource scenarios. ZNGIs are curved upwards or downwards when resources are complementary or antagonistic, respectively, and close to linear for the two weakly interacting scenarios. Within each scenario, the positions of the ZNGIs of the different species indicate where they lie along the generalist-specialist continuum: generalists have symmetrical ZNGIs at equal distance from the two resources axes while specialists on one resource have asymmetrical ZNGIs closer to that resource’s axes. The superimposed supply points help visualize environmental heterogeneity in each scenario. In the homogeneous case (top row), we also show how the consumption by the single species or pair of species (colored or gray arrows & points, respectively) that dominates after community assembly decreases resource availability from the supply point to the resource levels at equilibrium, which is the outcome of invasion analysis (See main text and Box 1). Similar arrows could have been used to show the variation in outcome from each supply point in the heterogeneous case (lower panels) but are omitted for clarity purposes.

We also considered how adding environmental heterogeneity at the local scale affects the results obtained from the Complementarity-Selection (Loreau and Hector 2001) and Community Assembly components methods. We ran our model assuming two different, spatially-implicit scenarios for the supply of resources over the local community: homogeneous resource supply and heterogeneous resource supply. The homogeneous environment was characterized by a constant influx of both resources (i.e. a single ‘supply point’, *sensu* Tilman 1982). The second scenario examines a heterogeneous environment, where species compete in an ecosystem characterized by 100 microhabitats, each with a separate supply point (see Fig. 1).

### Community Assembly Outcomes

To model the community assembly process, we viewed a BEF experiment as having the following sequence of events: a) there is a filter on the species pool (taken as a subset from the total species pool of the experiment), that is imposed by the researchers via selective seeding and/or weeding, b) interactions among the remaining species lead to some of the species going extinct, and c) there are density-dependent responses among the remaining species that then determine ecosystem attributes.

For each resource competition scenario (complementary vs. antagonistic, homogeneous vs. heterogeneous), we imitated these BEF conditions by generating multiple individual species pools with different subsets of species from the global pool that differ in their species richness and composition. In practice, we created a global pool of ten species whose acquisition traits for the two resources are regularly spaced along a generalist-specialist continuum defined by a trade-off between the two acquisition traits that parametrize the growth rate *g_j_*(*R*_1_, *R*_2_) (Appendix A). Our species pool of ten species yields 1023 possible subsets of this species pool, including the 10 different monocultures and the full species pool. We then considered the equilibrium states of communities initialized with all possible subsets of this global pool, arranged by their starting richness (e.g., 10 one-species communities, 45 two-species communities, etc.). This was performed using a global invasion analysis (Box 1), based on ZNGIs and their envelopes (Koffel et al. 2016). In the homogeneous case (a single supply point), only one or two species can subsist at equilibrium on two resources (Levin 1970) but more species can coexist if supply points are heterogeneous. In the heterogeneous case, 100 supply points were drawn from a multivariate normal distribution with the same mean as in the homogeneous case and a variance of 0.5 (See Fig. 1). The equilibrium community state was determined for each supply point. Because these equilibria for both the homogeneous and heterogeneous cases are deterministic, only one or two species will ultimately persist for any single supply point (more species can coexist for multiple supply points), and the identities of coexisting species and their abundances are predetermined by their growth functions and consumption vectors. Each resulting equilibrium community was characterized by the identity of the species remaining after community assembly and their equilibrium biomass. In the homogenous case, total ecosystem function is simply the sum of biomass for all species in the equilibrium community. For the heterogeneous case, equilibrium biomasses were obtained by averaging over the set of supply points, enabling a more direct comparison with the homogeneous case.

We then consider key properties of the communities that emerge under each resource scenario (across diversity levels and all initial compositions). These include the final biomass of each species, which together determine the total biomass or function of the ecosystem, and realized species richness (the final number of species present). This information for each of the subsets was then used in the Complementarity-Selection and Community Assembly calculations.

### Complementarity-Selection analysis

The outputs from our competition models serve as inputs to the Complementarity-Selection approach (Loreau and Hector 2001). For the purposes of this calculation, we assumed all species had equal ‘initial’ abundances. The decomposition divides total function into the Selection effect (SE) and Complementarity effect (CE) for each of the 1023 communities. SE and CE were averaged across all possible species pools at each level of introduced richness regardless of how many species actually existed at equilibrium.

### Community Assembly analysis

No supplementary assumption was needed to apply the Community Assembly components (Fox and Kerr 2012; Bannar-Martin et al. 2018). We compared the community initialized with the entire pool of ten species (our baseline, *sensu* Bannar-Martin et al. 2018) to each less diverse community. Using the Community Assembly configuration of the Price equation as described in Bannar-Martin et al. (2018), we calculated for each community the impact on ecosystem function of species lost (SL), species gained (SG) and the changes in function of shared species (CDE) relative to the final community obtained for the most diverse species pool. We then averaged each of these components across all possible species pools within each level of introduced richness.

### Comparison with experimental studies

To illustrate how our approach can provide insights about community assembly processes driving patterns of community richness and the functioning of ecosystems in empirical systems, we applied the same analyses to data from a well-studied, representative BEF experiment conducted at Jena (Roscher et al. 2004; 2005; Weigelt et al. 2016). Established in 2002, this experiment manipulated grassland communities, controlling the seeded species richness (2, 4, 6, 8, and 16 species, as well as the full 60 species pool) and functional group representation (1 to 4 groups) of 66 plots; additional small monocultures (3.5 x 3.5 m plots) were established for each species. Randomly selected subplots were sampled in the spring and summer of each year: aboveground biomass was clipped, sorted to species, dried, and weighed (see Roscher et al. 2004, 2005 for additional experimental details). Roscher et al. (2005) found evidence of positive selection and complementarity effects across plots with 2 - 8 species, focusing on data from May of the second year of the experiment (2003). Using data available from Weigelt et al. (2010), we expanded this Selection-Complementarity analysis to cover 2003-2008 (focusing on spring data, as do Roscher et al. 2005), and augmented it by applying the Community Assembly approach. Prior to analysis, we averaged species’ biomasses across replicate subplots within each experimental plot. The results provide an empirical case study for comparison with our modeling exercise.

Complementarity-Selection results for each year are based on monoculture biomass data collected during the corresponding year. Generally, monoculture biomasses declined over the six-year horizon of this analysis (Marquard et al. 2013; declines in polyculture yield were weaker). Consequently, there is a tendency for the magnitude of Complementarity-Selection effects to grow more pronounced over time, due to their mathematical definition (Loreau and Hector 2001). As the Community Assembly calculations are independent of monoculture data, they were not similarly affected.

## Results

In these initial analyses, we studied eight different scenarios with our model by varying the type of competition and the degree of resource heterogeneity as shown in Table S1 and Fig. 1. Different scenarios produced qualitatively distinct relationships between realized species richness (at equilibrium) and initial species richness (Fig. 2A), although ecosystem function (here total biomass or yield) was always a saturating function of initial species richness (size of the species pool; Fig. 2B), These results were similar for different combinations of resource type and/or environmental heterogeneity. Regarding our two approaches for analyzing communities, we focus on qualitatively distinct outcomes as follows:

**Figure 2.**
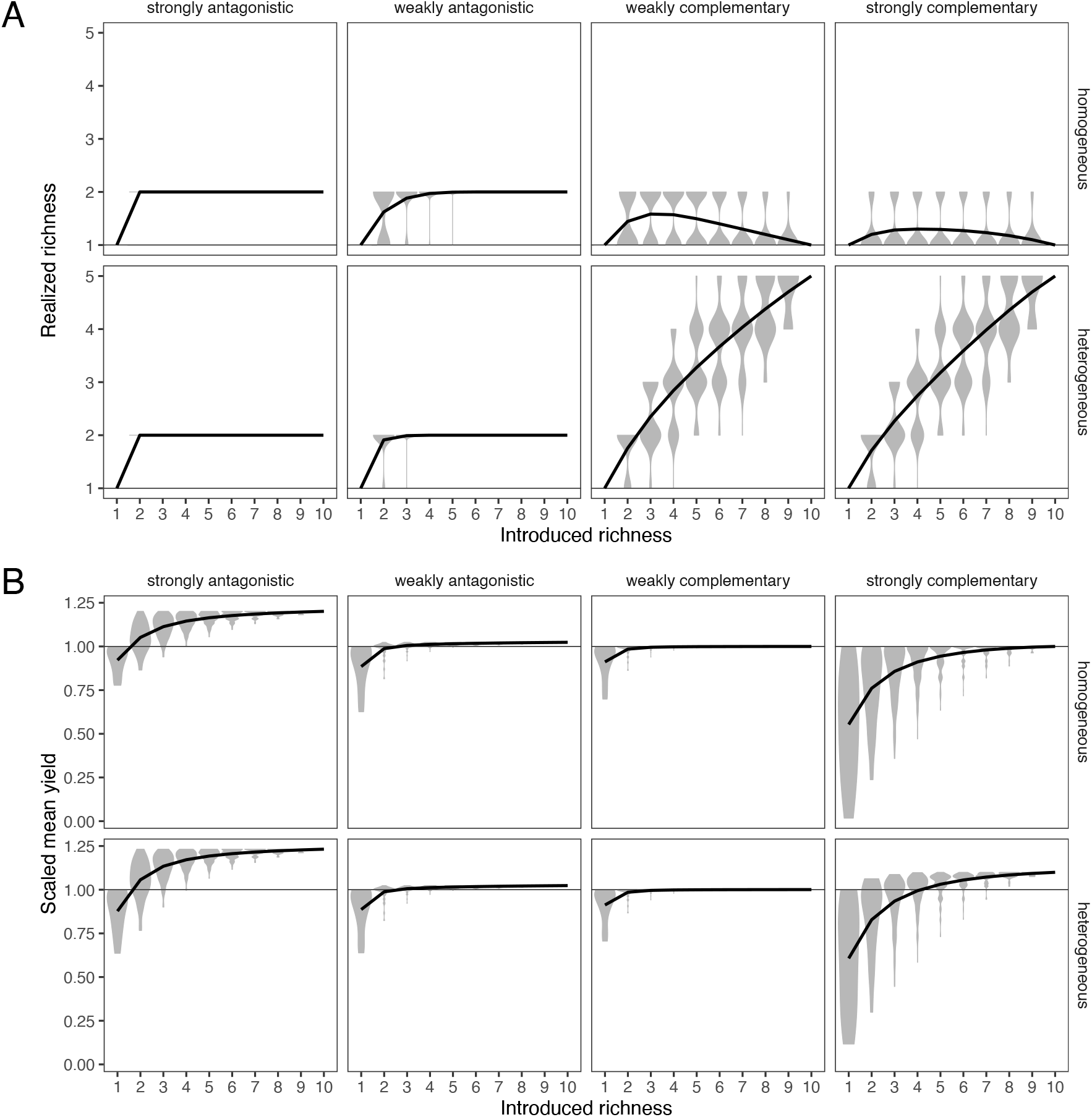
A) Realized richness obtained after species sorting (violin plots), averaged across all possible species pools at each level of introduced richness (solid black line). In the four antagonistic resource scenarios, realized richness quickly saturated at two species. In the complementary resource scenarios, average realized richness is hump-shaped in the homogeneous case, but increasing in the heterogeneous case. B) Total community yield obtained after species sorting (violin plots), averaged across all possible species pools at each level of introduced richness (solid black line); values are scaled relative to the yield of the most productive monoculture (horizontal reference line). All scenarios give saturating responses but in scenarios where resources are close to substitutable (central panels), moving from monocultures to polycultures causes weaker increases in yields.

### Antagonistic resources

Antagonistic resources, whether weak or strong, or supplied uniformly or heterogeneously, produced similar results. First, the maximum yield of polycultures was always greater than the maximum monoculture. Second the realized richness was always two species or fewer. In these scenarios, the average complementarity effect (CE) was always positive — setting aside the weakly antagonistic case with introduced richness of two — whereas the average selection effect (SE) was always negative (Fig. 3A). Finally, the average Community Assembly components always showed positive species gains (SG) and negative species loss (SL) components, whereas the CDE was always small or zero (Fig. 3B).

**Figure 3.**
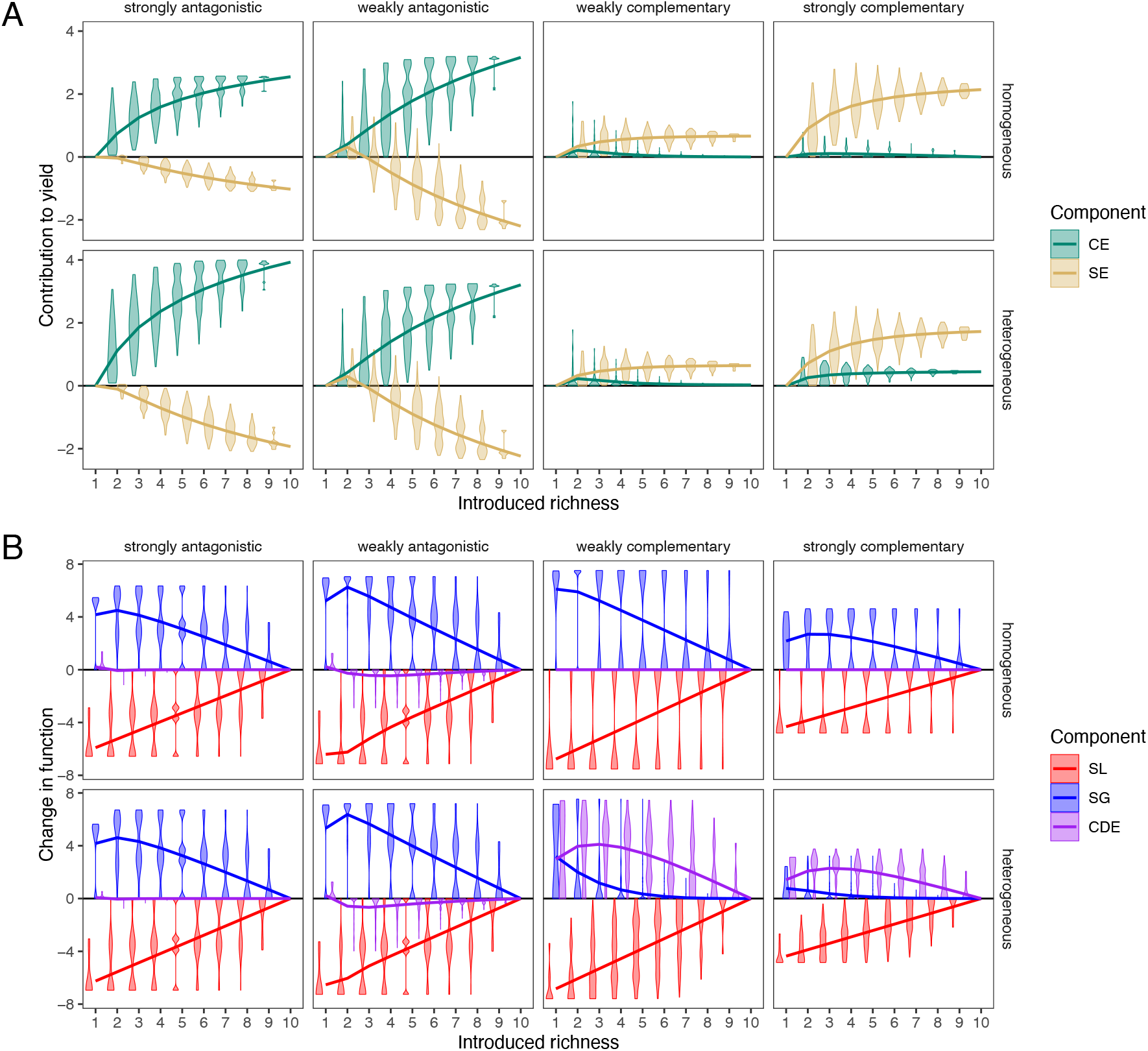
A) Complementarity (CE) and selection effects (SE) vary in magnitude and sign as a function of introduced richness, competitive scenario (columns), and supply point heterogeneity (rows). Antagonistic resources lead to CE > 0 and SE < 0, while complementary resources result in CE > 0 and SE > 0. B) Community Assembly components method, comparing the most diverse polyculture (initial richness of 10) against less diverse mixtures. CDE is nearly always zero or slightly negative, except when resources are complementary and have a heterogeneous supply. For both A) and B), violin plots and solid lines respectively indicate variability and average values across all possible species pools at each level of introduced richness (as in Fig. 2). Violins are jittered slightly horizontally for clarity.

### Strongly complementary resources in homogeneous environment

In contrast to antagonistic resources, results varied substantially depending on just how strongly resources are complementary and on the degree of environmental heterogeneity. Starting with strongly complementary resources (almost equivalent to essential resources), we found that maximum yield in polycultures did not differ from that of the most productive monoculture. Both selection (SE) and complementarity effects (CE) were positive, in contrast with antagonistic resources (Fig. 3A). The increase in yield with diversity was largely due to an increase in the mean selection effect: as the size of species pools grew, more productive competitors were more likely to be present. The relationship with complementarity was unimodal and reflected the increased role of complementarity on yield in small species pools where the better competitors were absent. The effects of complementarity were also generally small. Community Assembly components showed that effects of CDE were zero and that all of the effects involved positive effects of SG and negative effects of SL (Fig. 3B). We also found that mean realized species richness was a unimodal function of the size of the initial species pool in the homogeneous scenario (but still limited to two or fewer species; Fig. 2.A and S1). This means that the relationship between realized diversity and yield appears complex (Figure S1).

### Strongly complementary resources in heterogeneous environments

The patterns differed from the homogenous environment results in several respects. First, the maximum productivity was typically higher than that of the most productive monoculture. Also, realized species richness was now positively related to the size of the species pool. In contrast with the homogeneous case, the positive complementarity effect (CE) persisted at high diversity levels (Fig. 3A), while species gains (SG) had much smaller effects and CDE was positive, counteracting a similar SL effect (Fig. 3B). We also examined an intermediate scenario, with smaller variance in supply point heterogeneity, and found intermediate patterns between the homogenous case and the full variance case: intermediate SE and CE, and CDE and SG of comparable magnitudes (not shown).

### Weakly complementary resources

With resources that were only weakly complementary, the increase in yield with diversity was less pronounced. Both CE and SE results were similar to the complementary case (Fig. 3A), except that environmental heterogeneity did not support a sustained positive CE effect at high diversity levels.

### Relations between realized species richness and biomass

In each of our scenarios, we also examined how total biomass was related to the realized species richness (the number of species coexisting at equilibrium). Over all initial richness levels, there was generally a positive relationship between standing biomass and realized species richness (Fig. S1). However, scenarios differed in the strength of this relationship and in the structure of covariation between biomass and realized species richness when the size of the species pool was accounted for. In most scenarios (scenarios 1-6 and 8 in Table S1), we found that biomass was positively related to realized species richness and that this was true both at the global scale of analysis and within subsets of communities that differed in the size of the introduced species pool. In contrast, when resources were strongly complementary and environmental heterogeneity low (scenario 7), total biomass was negatively related to realized species richness for any given level of initial richness (Fig. S2).

### Mechanisms underlying relationships between competition scenarios and Complementary-Selection and Community Assembly patterns

The striking differences between the antagonistic and complementary resource competition scenarios captured by the Community Assembly components can be linked to the influence of resource type on the outcomes of community assembly. In the antagonistic case, a pair of extreme specialists dominates all the other species for any supply point (Box 1, panels A, B). This strongly affects both the complementarity and selection effects (CE and SE), and the Community Assembly components (SL, SG, and CDE). Dominance by a pair of specialists leads to a realized richness that cannot exceed two even in the heterogeneous case. In monoculture, each specialist strategy generally performs worse than the intermediate generalist strategies, which explains why the selection effect is negative. Yet, the two specialists together perform better than any other set of one or two species, driving the positive CE pattern. When one of the two dominant specialist strategies is absent (causing a negative SL component), it is replaced by a less specialized species with lower function than the missing specialist (which results in a positive SG component but |SL| > |SG|). The less specialized species competes more strongly with the remaining specialist due to greater niche overlap, reducing the performance of the remaining specialist, which creates the negative CDE.

In the complementary cases, there is usually a unique generalist species from the full species pool that dominates for a given supply point (Box 1, panels C, D). This dominant species is also the one that performs the best – better than any other set of one or two species – for this supply point, which explains why selection is positive and complementarity is weak and hump-shaped. The CDE of exactly zero in the homogeneous case is a direct consequence of having a single dominant strategy: either this dominant strategy is absent from a low diversity community, which will then ultimately share no species with the high diversity community, or the dominant strategy is present, producing an endpoint identical to the high diversity community. Finally, the positive CDE in the heterogeneous case arises because in the absence of one or more of the dominant strategies, other dominant species that still occur compensate for functional losses. Such CDE is larger than the SG component on average because the functional contributions of new species (able to join a community due to the loss of at least one dominant species) are smaller than increases in function occurring through compensation by the remaining dominants.

#### Box 1: Mechanisms

In this box, we give a mechanistic explanation of how different competitive scenarios determine the identity and function of persistent species. In turn, these will determine the sign and magnitude of the Community Assembly components that arise from comparing communities assembled from high and low diversity species pools. We consider both strongly antagonistic and complementary resource scenarios. We show a simplified example where the full species pool only includes five species, three of which appear in the low diversity subset. We focus on a homogenous environment (with one supply point; black dot) in the antagonistic example and a heterogeneous environment (with three supply points) in the complementary case.

These situations are represented and analyzed using the graphical approach to resource competition (MacArthur 1972, Tilman 1982, Leibold 1995, Chase and Leibold 2003, Koffel et al. 2016). We perform the invasion analysis by drawing the ZNGIs of the species in the community all together. The species whose ZNGIs are entirely located above other species’ ZNGIs are systematically excluded (e.g. purple, orange, and yellow in A). When the impact vectors of the remaining species (red and blue in A) are combined with a supply point, we obtain the set of these species that actually end up persisting in the equilibrium community (e.g. red and blue coexist in A). In a heterogeneous environment (C and D), we perform the invasion analysis for each supply point independently and average the productivity of the resulting persisting species to obtain the equilibrium community.

In the antagonistic case, when all members of the full species pool are initially present (A), the three intermediate species (purple, orange, yellow; dotted lines) are excluded by the two extreme specialists (red and blue; solid lines), which coexist. However, if one of these two specialists (e.g., blue) is randomly absent from a low-diversity community (B), it is replaced by its closest functional equivalent among the intermediate species (here, yellow). This species is not completely specialized on resource 2, so it does not perform as well as the lost specialist. Due to weak competition with this intermediate species, the performance of the remaining specialist (red) is lower than in the full species pool situation.

In the complementary case, the most diverse species pool (C) experiences the exclusion of the two most extreme species (orange and yellow), while the three intermediate species (blue, red, and purple) coexist, as each one is locally adapted to a particular supply point. If two of these intermediate species, which dominate the full species pool, are absent in a smaller community (e.g., red and purple), an extreme species (e.g., yellow) can survive and coexist with the remaining dominant species (blue). This extreme species (yellow) only partially compensates for the loss of function that occurs due to the absence of the purple species. Similarly, the remaining dominant species (blue) partially compensates for the loss of function due to the absence of the red species by increasing in abundance.

**Figure Box 1:**
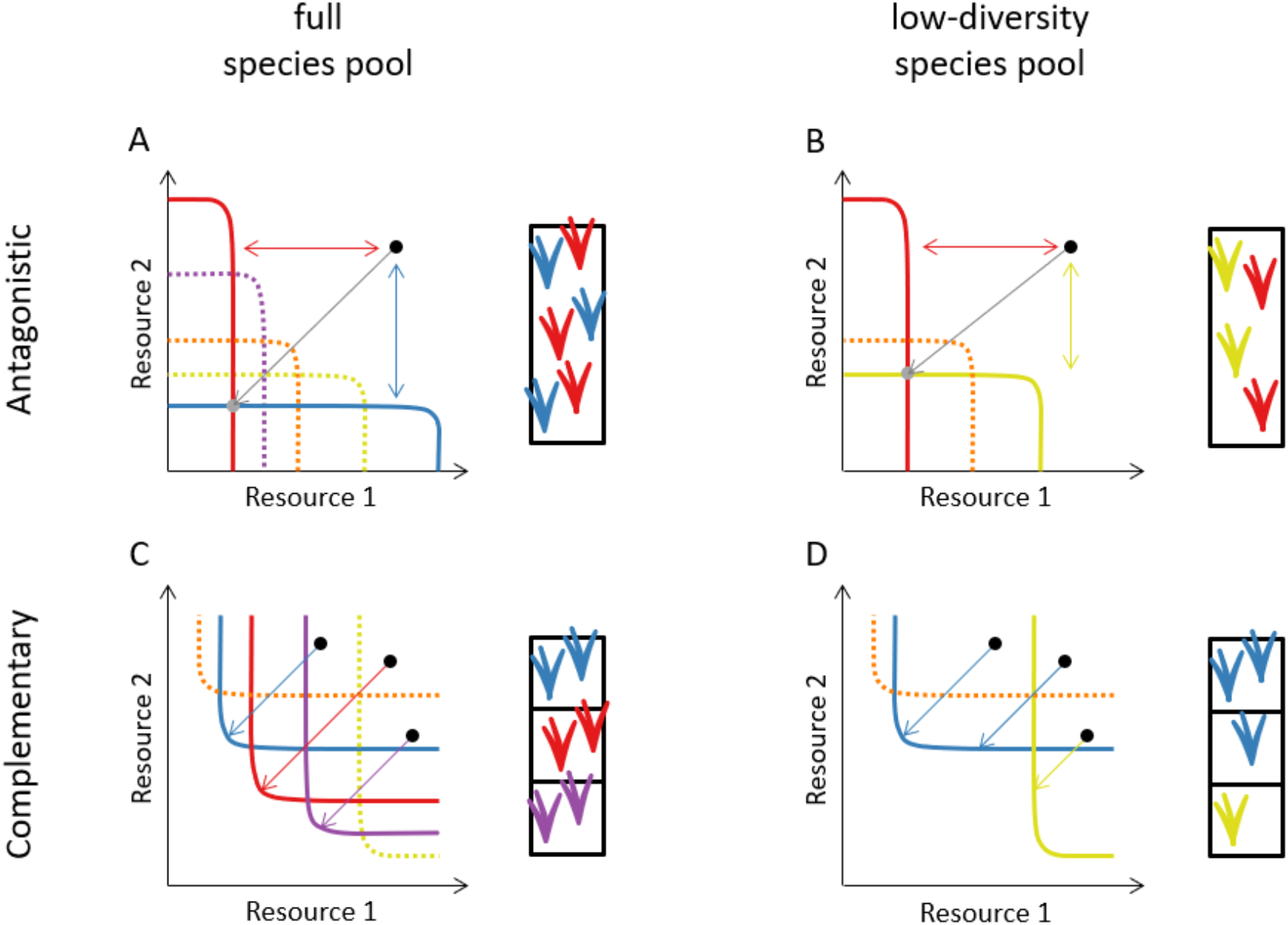

### Comparisons with experimental data

Although most of our scenarios (with the exception of scenario 7) produce broadly saturating BEF patterns, our results indicate that we can qualitatively discriminate among most of these eight different community assembly scenarios (Table S1) by combining the Community Assembly (Fox and Kerr 2012, Bannar-Martin et al. 2018) and the Complementarity-Selection (Loreau and Hector 2001) analyses. This suggests that applying these methods to empirical data may offer insights into which competitive scenarios drive assembly and give rise to empirically-obtained BEF relationships (or at least provide evidence about which mechanisms may be ruled out). To assess this, we analyzed data from a representative BEF experiment, conducted at Jena, Germany, to determine if we could identify the resource competition mechanisms underpinning the functioning of this grassland ecosystem. We used six years of data from the Jena experiment, starting in 2003 (a year after the experiment was established), to consider whether results were stable over time.

We found that the data from Jena did not match any of the scenarios shown in Table S1. On the one hand, we see a divergence between increasingly negative selection effects and increasingly positive complementarity effects from 2003 to 2008 (Fig. 4), creating a pattern consistent with the antagonistic resource scenario using the Complementarity-Selection approach (Fig. 3A, see also Marquard et al. 2009). On the other hand, we see persistent trends in the Community Assembly components, including positive CDE effects across the entire time span (Fig. 4), which were only observed in our model scenarios given competition for heterogeneous, complementary resources (Fig. 3B).

**Figure 4.**
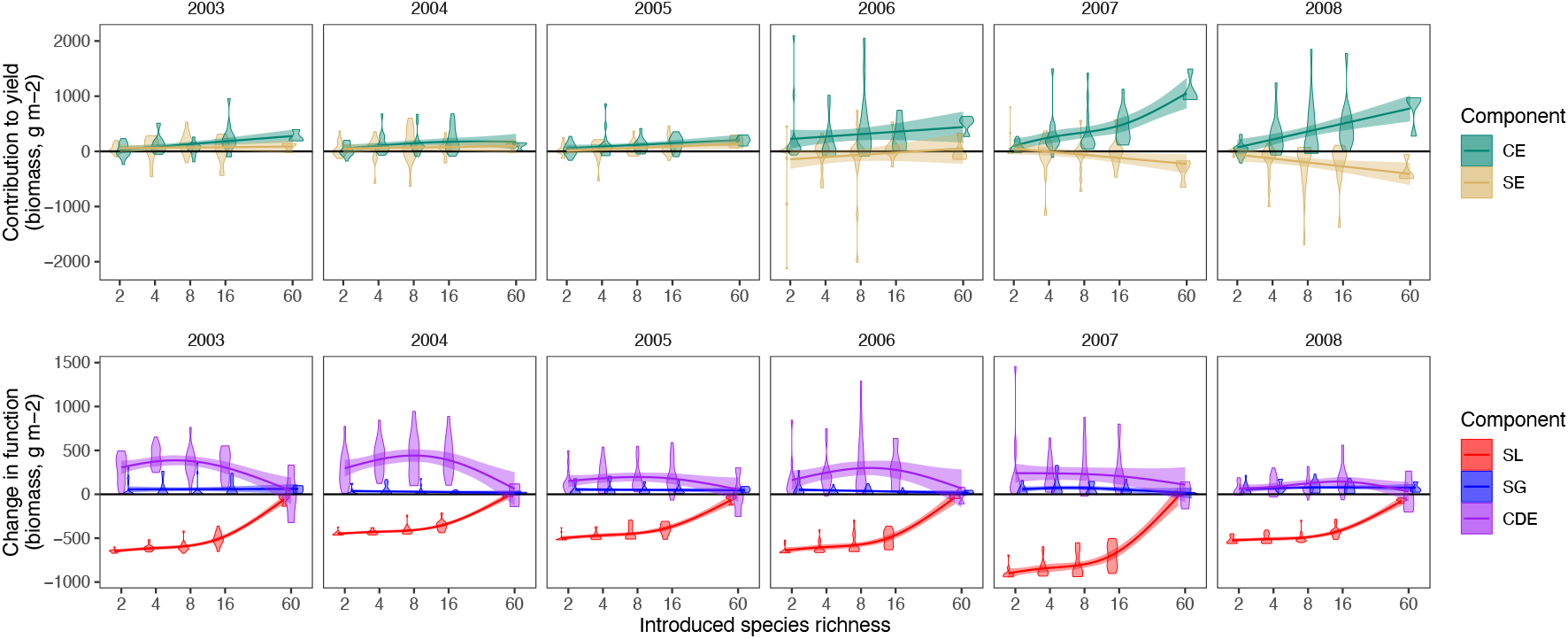
Complementarity (CE) and selection (SE) effects (top row), and Community Assembly components comparing the most diverse (60 species) community to less diverse communities (bottom row) over successive years at the Jena Experiment, as a function of introduced species richness (log scale). Early in the experiment, CE and SE were both positive, and weakly related to diversity (as found by Roscher et al. 2005). However, by the end of the period we examined, the CE effect was positive and increased strongly with diversity, while the SE effect was negative and decreased with diversity. Although there is some variation in the magnitude of the Community Assembly components through time, at all lower diversity levels, CDE effects are consistently positive, SG is negligible, and SL strongly contributes to declines in function. Trend lines show Generalized Additive Models (GAMs) fit to the observed plot-level values, including confidence bands; violins were jittered horizontally for visual clarity.

Consequently, the analysis of Complementarity-Selection effects (Fig. 4) is consistent with antagonistic resources, be it in homogeneous or heterogeneous supply points (compare with Fig. 3A), whereas the Community Assembly components suggest a scenario of complementary resources in the presence of environmental heterogeneity (comparing Fig. 4 with Fig. 3B). This indicates, perhaps unsurprisingly, that the eight simplistic scenarios we considered probably lack additional mechanisms involved in regulating or modifying the effects of community assembly on ecosystem function (and hence the metrics used to study it).

As a result, we considered other possible factors that might have affected community assembly at Jena. One of these is that the empirically observed dynamics may not have reached steady state as assumed by our models above, a hypothesis supported by the directional changes in Complementarity-Selection effects across the first three to four years of the experiment (Fig. 4). To address this possibility, we simulated the temporal dynamics of community assembly for the heterogeneous, complementary resources scenario (because previous work has mostly emphasized that this scenario is more likely than others for plants; see Tilman 1982, Tilman et al. 1997, Gross and Cardinale 2007, Hodapp et al. 2016). Starting from an initial condition where all species are seeded at an equally low density, we let their populations grow and compete over time. When a species’ density dropped under a certain minimal threshold (set equal to 10^-4^), we considered it extinct and set its density equal to zero. We then extracted species densities at representative time points and applied both partitions as previously described. Although this exercise shows that there can be important transient dynamics in the community assembly dynamics of BEF experiments (Fig. S4), none of the resulting patterns are consistent with the patterns we observed in Jena (Fig. 4). In fact, the qualitatively stable distribution of metric values for Jena data after the fourth year suggest that the empirical system was near, if not at, equilibrium. Intriguingly, we found that the dynamics in our model (Fig. S4) showed a strong qualitative shift in the relative importance of CE and SE through time and a somewhat subtler shift from linear to unimodal CDE and an increase in SG through time. However, the data from Jena show no such trends.

We also considered that facilitation, perhaps due to the nitrogen (N) fixing effects of legumes, may play an important role in assembly, and consequently affect ecosystem functioning. To address this, we adapted the model of Koffel et al. (2018) that combines strongly complementary resource competition with facilitation, and again applied Complementarity-Selection and Community Assembly analyses (see Appendix B). With high environmental heterogeneity and strong facilitation under high initial N-limitation, our results (Fig. 5) correspond at least qualitatively to our observations at Jena. Both the facilitation model and Jena data demonstrate greater complementarity effects (CE) in comparison to selection (SE) and, at least for Jena’s 2007 and 2008 data, a negative (Compare Fig. 4 and Fig. 5A). This seems consistent with observations from the Jena experiment that showed that N fixation rates by legumes, and thus potentially facilitation, increased from 2004 to 2008 (Roscher et al. 2011). The Community Assembly components also show similarities, with a substantial and humped CDE effect that emerges from the model (Fig. 5B) also being seen in the Jena data (Fig. 4). The addition of facilitation creates a combination of Complementarity-Selection effects and Community Assembly components patterns that otherwise appear to point to divergent resource competition models in the absence of facilitation.

**Figure 5:**
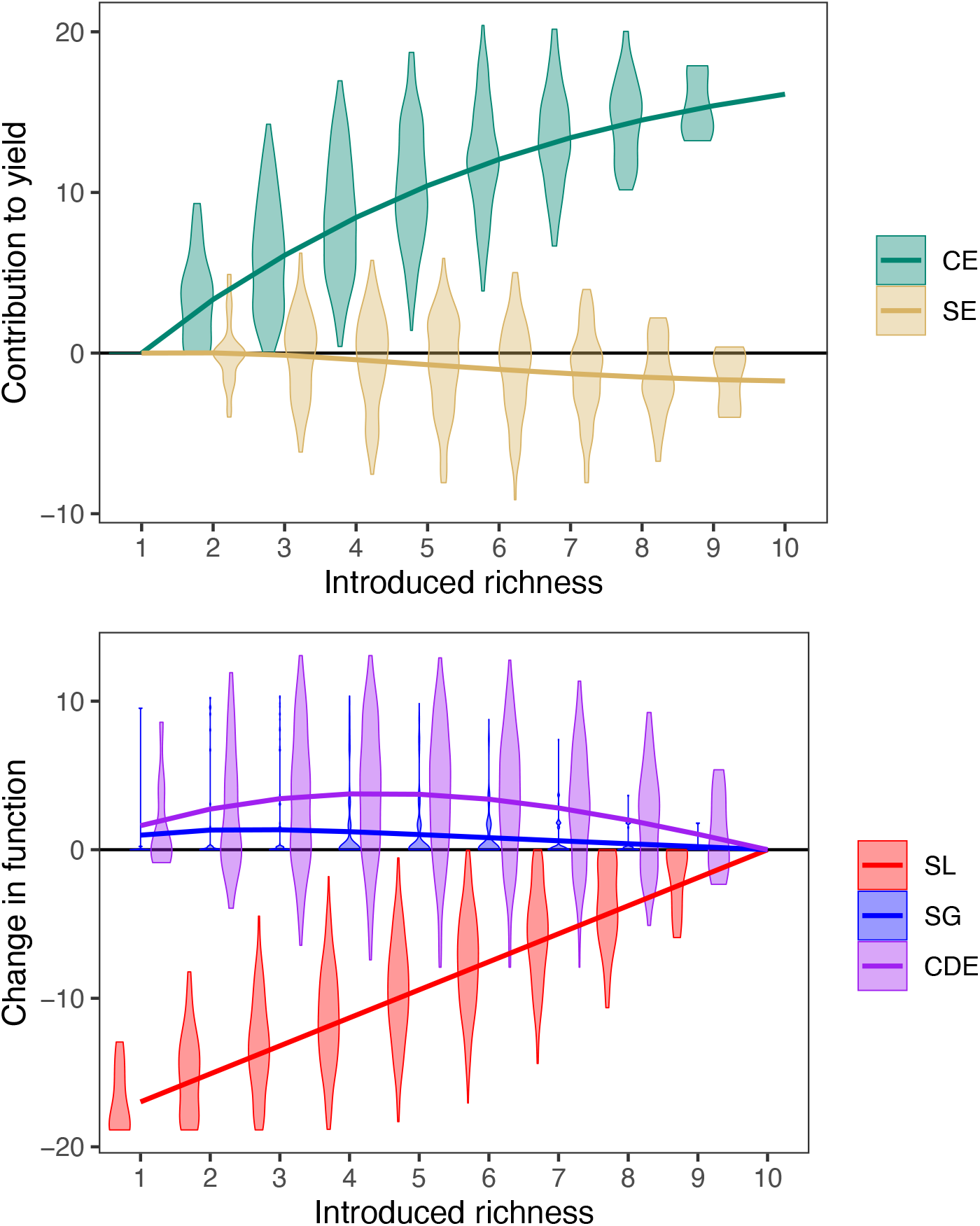
Model results after including nitrogen fixing plants that compete with non-fixers, including Complementarity-Selection effects (top) and Community Assembly components (bottom). Violin plots and solid lines respectively indicate variability and average values across all possible species pools at each level of introduced richness (as in Fig. 2).

To evaluate this hypothesis, we conducted separate Complementarity-Selection and Community Assembly analyses for Jena plots that did and did not include N-fixing plants (legumes) in their species pools (Fig. 6). Plots that contained legumes showed results that qualitatively match the facilitation scenario (compare with Fig. 5) and differed from plots without legumes as might have been predicted. However, plots that lacked legumes exhibit patterns that are both qualitatively inconsistent with any of our scenarios and relatively weak, perhaps due to subtler mechanisms. Further exploration of these effects is beyond the scope of the present publication.

**Figure 6:**
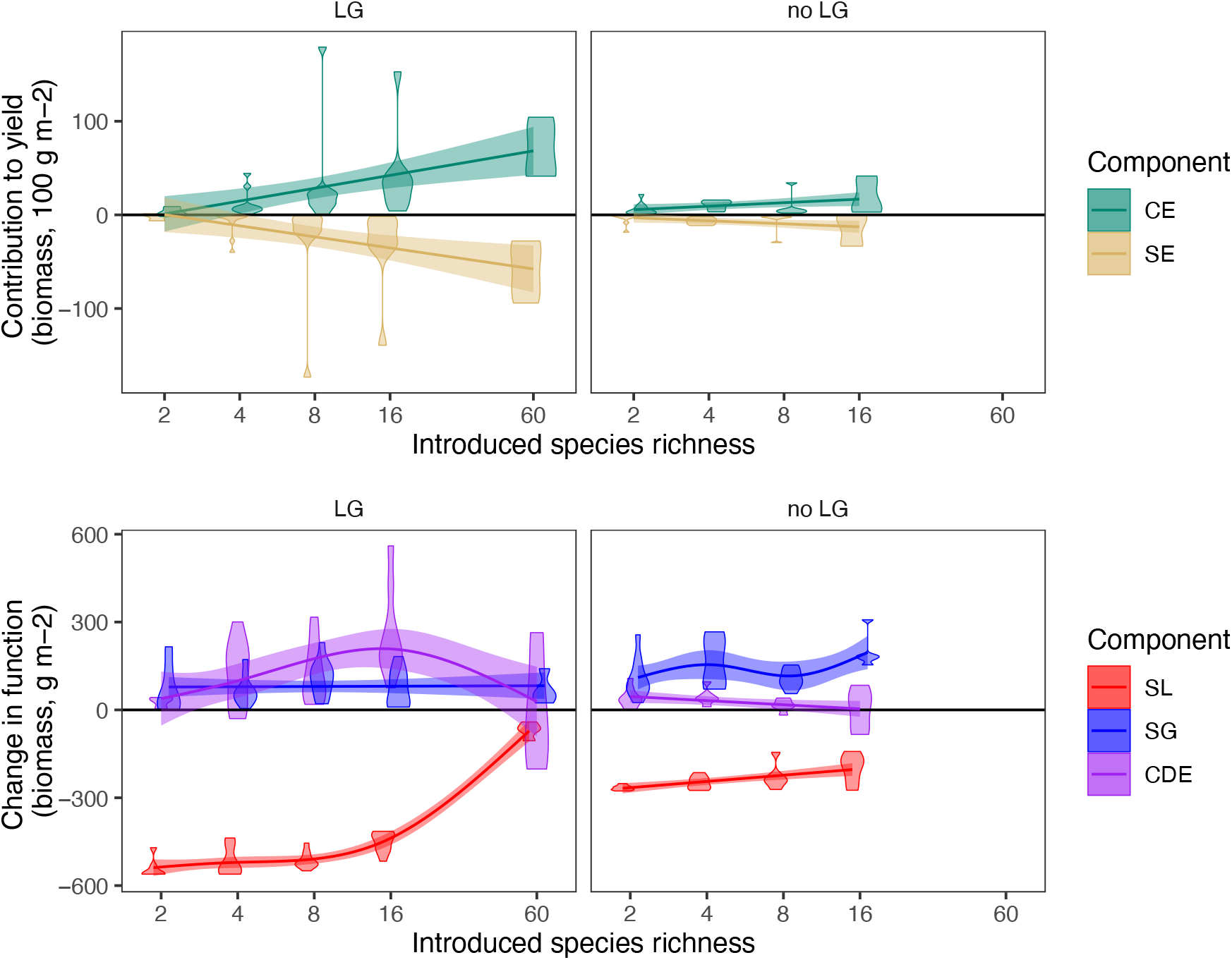
Complementarity-Selection effects (upper panels) and Community Assembly components (lower panels) for subsets of the Jena experiment that contained at least one legume (LG, left panels) or none (right panels). Trend lines show Generalized Additive Models (GAMs) fit to the observed plot-level values, including confidence bands.

## Discussion

There are two major implications of our work. First, we find that the joint use of the Community Assembly (Bannar-Martin et al. 2018) and the Complementarity-Selection (Loreau and Hector 2001) approaches can be a powerful tool in inferring (or excluding) mechanisms that drive community dynamics and consequently the functioning of ecosystems. Second, we find that applying this joint approach to a real-world system (the Jena BEF experiment) enabled us to rule out a series of purely competitive community mechanisms – where using only a single approach would have led us to inaccurate conclusions. This ultimately resulted in new hypotheses, and the conclusion that facilitation by N-fixing plants is critical. This suggests that the joint evaluation of Complementarity-Selection effects and Community Assembly components could be a powerful tool for identifying plausible mechanisms that relate community assembly to the functioning of ecosystems.

### Validating the joint use of Complementarity-Selection and Community Assembly approaches to interpret mechanisms underlying patterns in BEF studies

Numerous studies in controlled and replicated conditions (such as those that prevail in the large BEF literature) have shown that there is generally a positive and directional effect between the size of the species pool with access to a community and the function of at least some aspects of ecosystems (especially total biomass). Previous work on relationships between species pool diversity and ecosystem function has largely focused on two very general effects that can explain this pattern: selection (the species pool, when large, is more likely to include highly competitive and highly productive species) and complementarity (the species pool, when large, is more likely to include species that coexist and whose production is less mutually inhibitory). To date, theoretical explanations for these two general effects have focused on resource competition and used models based on complementary resources (e.g., Tilman et al. 1997, Gross and Cardinale 2007, Leibold et al. 2017).

However, other competitive mechanisms may be involved, and there has been little consideration of whether or not they would impart distinct complementarity and selection effects. Our results show that indeed different mechanisms can produce similar patterns of complementarity and selection, as the antagonistic resource and facilitation scenarios both lead to increasing positive complementary effects (CE) and decreasing negative selection effects (SE) with increased diversity (compare Fig. 3A with Fig. 5). This means that attributing specific Complementarity-Selection patterns to particular mechanisms may fall short. However, added insights from the Community Assembly approach (Bannar-Martin et al. 2018) improved the situation, as these two scenarios differed by the sign of the CDE and its relative magnitude compared to SG. Together, the joint use of Community Assembly and Complementarity-Selection approaches provided well-defined criteria by which we could distinguish between competition for complementary vs. antagonistic resources as well as evaluate the role of environmental heterogeneity and facilitation.

### Challenges identifying mechanisms that drive community-ecosystem function relations

Our study also reiterates the fact that we do not yet understand all the mechanisms that drive community-ecosystem relationships. Patterns in the data from the Jena experiment did not correspond to any of the eight scenarios we initially modeled, which correspond to the classic scenarios of resource competition in community ecology, and allowed us to reject all eight of these hypotheses. On the one hand, this shows how powerful the joint application of Community Assembly and Complementarity-Selection metrics can be. However, it also shows that the mechanisms governing the diversity and function of communities must involve other processes besides simple resource competition. This is not totally surprising since our theoretical models are caricatures of how species interact in communities, where other factors including spatial structure, grazers, pathogens, and mutualists may matter.

Further progress in this arena may require considering some of these other scenarios, aided by Community Assembly and Complementarity-Selection approaches. For example, we illustrated this by considering (1) non-equilibrium dynamics and (2) facilitation by an N-fixer; the latter scenario qualitatively agreed with observed patterns at Jena where other scenarios failed. This provides a new hypothesis, which may be address by subsequent research – a critical step, as we note that a correspondence between theory and observation is not definitive proof that a particular mechanism is operating. For example, our model of facilitation was based on N-fixation, but results similar to ours could arise from a broader range of facilitation mechanisms at play in BEF experiments, perhaps related to other abiotic factors (Wright et al. 2017) or based on soil organisms (Eisenhauer et al. 2012). Alternatively, other mechanisms besides facilitation might be capable of generating similar patterns. Such additional mechanisms may play a secondary role as revealed in the analysis of plots that did not contain legumes as shown in Fig. 6. Ultimately, for Complementarity-Selection and Community Assembly approaches to advance science, any given analysis and its results needs to be followed up with additional research and experimentation to verify putative mechanisms.

### Community assembly and the functioning of ecosystems outside of BEF designs

The design of BEF experiments imposes constraints on the assembly of communities that may not be present in real ecosystems. For example, species invasions, disturbance and dispersal dynamics, environmental change, and stochastic processes are all common in real communities (e.g. Thompson and Gonzalez 2016, Hodapp et al. 2016, Leibold et al. 2017, Bannar-Martin et al. 2018). We have shown that Complementarity-Selection and Community Assembly approaches offer useful insights when applied to BEF-style scenarios and data that limit these kinds of effects. This utility may extend to communities (and data) that differ from BEF designs, but requires careful thinking.

In particular, the Complementarity-Selection approach requires estimates of ecosystem function in monocultures. These are readily available in most ‘classic’ BEF experiments as a *pro-forma* aspect of their design. However, these are almost always absent in other experiments that examine how community assembly affects ecosystem functions (e.g. disturbance, invasions, extinctions, environmental change), let alone in observational data. The logistical challenge is even greater in studies examining the effects of environmental change on ecosystem function, as monoculture treatments would be required for all the relevant environmental conditions. Additionally, many organisms are not easily grown in monocultures due, for example, to dependencies on biotic resources, mutualists, or movement. Thus, the scenarios where the Complementarity-Selection approach can be used are limited.

In contrast, the Community Assembly approach can be calculated for any pair of communities, without monoculture data. Given the design of BEF studies, we opted to use the most diverse species pool as our baseline for comparisons, but this is not required (see Bannar-Martin et al. 2018 for examples). While the Complementarity-Selection approach is constrained by the requirement of monoculture data, it did appear to more clearly distinguish between antagonistic and complementary resource scenarios. This ability in part may rest in the definition of these metrics: both complementarity (CE) and selection (SE) are affected by modest changes in the abundance of a species. In contrast, of the three Community Assembly components, SL and SG are mostly influenced by the actual loss or gain of species from a community; all effects of less extreme changes in abundance are relegated to the CDE component alone. For this reason, while it is possible to apply a Community Assembly approach in a manner directly analogous to Complementarity-Selection (contrasting an observed polyculture with an idealized polyculture where species perform proportional to their monocultures), the results are somewhat less informative.

Our work shows that, while both methods exhibit their own advantages and limitations, together they provide a more powerful means of diagnosing mechanisms. Future work could focus on continuing to refine these metrics. This might include developing analogs of Complementarity-Selection partitions that do not require monoculture data. Alternatively, new or existing extensions of the Community Assembly approach (e.g., 5-part Price equation; Fox & Kerr 2012; Bannar-Martin et al. 2018) could be applied to these or other model scenarios, perhaps focusing on further dissecting the CDE component. For example, while empirical applications of the trait-based version of the Price equation (Fox & Harpole 2008) can be hampered by the availability of trait data, in modeling exercises these traits are explicitly known (e.g., species’ ZNGIs in our scenarios). Further work could elucidate not only the mechanisms structuring ecosystem function across communities of different diversity levels but relate ecosystem properties to the functional composition of communities.

### Implications for future work

These encouraging results suggest a number of ways whereby the analysis of community assembly and its effects on ecosystem function could be improved:

a. Most obviously, one could continue to propose and test other mechanisms, such as plant-soil feedbacks and multi-trophic interactions, that could have produced the patterns we observed in the Jena data, eventually potentially creating a ‘periodic table’ for how the Complementarity-Selection and Community Assembly metrics map onto a diverse array of community assembly scenarios. While there is some allure to this suggestion, there are a considerable number of interacting mechanisms to test requiring careful thinking about how to construct and interpret models (e.g., considering transient dynamics), or how to parse varying contrasts in the data (e.g., considering facilitation by legumes).
b. Discriminating among the wide array of mechanisms driving community assembly and its effects on ecosystem function likely requires moving beyond simple dichotomous tests with one or two metrics since many alternative models/mechanisms are capable of reproducing low-dimensional patterns. Here we demonstrated how the addition of several carefully considered metrics could enhance our understanding of these mechanisms, as well as our ability to differentiate them. It is worth considering whether other additional metrics could similarly advance understanding. The degree to which this is so would depend on finding metrics that reveal distinct aspects of mechanisms and that, thus, are not simply redundant with each other. It is also useful when the link between mechanism and metrics is relatively transparent (as we explored in Box 1). More complex relationships between mechanisms and metrics might require basic modeling or experimental studies to help interpret the results. The increase in the number of diagnostic metrics, however, may be necessary if theoretical models increase in complexity by including missing processes or by assessing different assumptions to increase inference resolution (e.g. Cabral et al. 2019).
c. Another reason to consider additional metrics is that not all data will be suitable for the same metrics. In our case, while the Complementarity-Selection approach was particularly useful in discriminating between different types of resource competition, the approach is constrained by the need for monoculture data. We will need new techniques if we wish to apply these types of approaches to a wider array of experimental conditions, with designs that do not closely target BEF relations or that involve organisms where such monocultures are logistically difficult (e.g. birds, migrating organisms, species-rich tropical communities).
d. While we need to explore other metrics to use in studies other than BEF experiments, it would still be interesting to apply the methods in this paper to other BEF experiments. We suspect that this may likely show substantial variation in results across locations, habitats, and organisms. For example, it would be interesting to compare results at Jena with data from sites that are not as likely to be nitrogen limited. Adequately considering such variation would require a far more comprehensive theoretical and analytical effort than we could undertake in this initial paper.
e. Even more challenging is considering how the size of the species pool jointly affects biodiversity, community composition, and ecosystem functioning under other scenarios than those at play in classical BEF experiments. Numerous challenges to BEF research have focused on the degree to which BEF experiments may be unrealistic representations of how biodiversity and ecosystem function are likely to interrelate because the experimental setup for community assembly in BEF experiments is so artificial (Wardle 2016). We believe the Community Assembly approach can shed light on other situations including arrivals and removals of invasive species as well as environmental change (see Bannar-Martin et al. 2018, Kremer et al. in prep). Linking these scenarios to explicit theoretical models (as we did here by asking how resource types and environmental heterogeneity affect a BEF experiment) represents an exciting avenue for future devoted work.

## Software availability

An R package is available to facilitate future work (including analyzing other biodiversity experiments) https://github.com/ctkremer/priceTools

## Acknowledgements

This paper is a joint effort of the working group sCAFE and an outcome of a workshop supported by sDiv, the Synthesis Centre of the German Centre for Integrative Biodiversity Research (iDiv) Halle-Jena-Leipzig (DFG FZT 118). The Jena Experiment is funded by the German Research Foundation (FOR456/1451) with additional support from the University of Jena and the Max Planck Society. MAL was also supported by a research award from the Alexander Von Humboldt Foundation.

