## Supplementary information for "Modeling How Community Assembly Alters the Functioning of Ecosystems"

**Table S1**. The eight modeling scenarios in our study, which differ in the type of resource competition and degree of environmental heterogeneity they feature, as described in the text and illustrated in Fig. 1. Our results showed that both competition and heterogeneity produced variable effects on complementarity (CE), selection (SE), species loss (SL), species gain (SG) and context dependence (CDE). We also found that comparisons between SG and CDE were informative. The magnitude of these effects is summarized qualitatively through the categories “exactly zero” (0), “approximately zero” (≈ 0), “negative” (−), “positive” (+) and “strongly positive” (++).

| Scenario | Resource type | Environmental heterogeneity | Predicted effect (see Results) on: | | | | | |
|---|---|---|---|---|---|---|---|---|
| | | | CE | SE | SL | SG | SG vs CDE | CDE |
| 1 | strongly antagonistic | homogeneous | ++ | − | − | ++ | > | ≈ 0 |
| 2 | strongly antagonistic | heterogeneous | ++ | − | − | ++ | > | ≈ 0 |
| 3 | weakly antagonistic | homogeneous | ++ | − | − | ++ | > | − |
| 4 | weakly antagonistic | heterogeneous | ++ | − | − | ++ | > | − |

| | | | | | | | | |
|---|---|---|---|---|---|---|---|---|
| 5 | weakly complementary | homogeneous | ≈ 0 | + | − | ++ | > | 0 |
| 6 | weakly complementary | heterogeneous | ≈ 0 | + | − | + | < | + |
| 7 | strongly complementary | homogeneous | ≈ 0 | + | − | ++ | > | 0 |
| 8 | Strongly complementary | heterogeneous | + | + | − | + | < | + |

### Figures

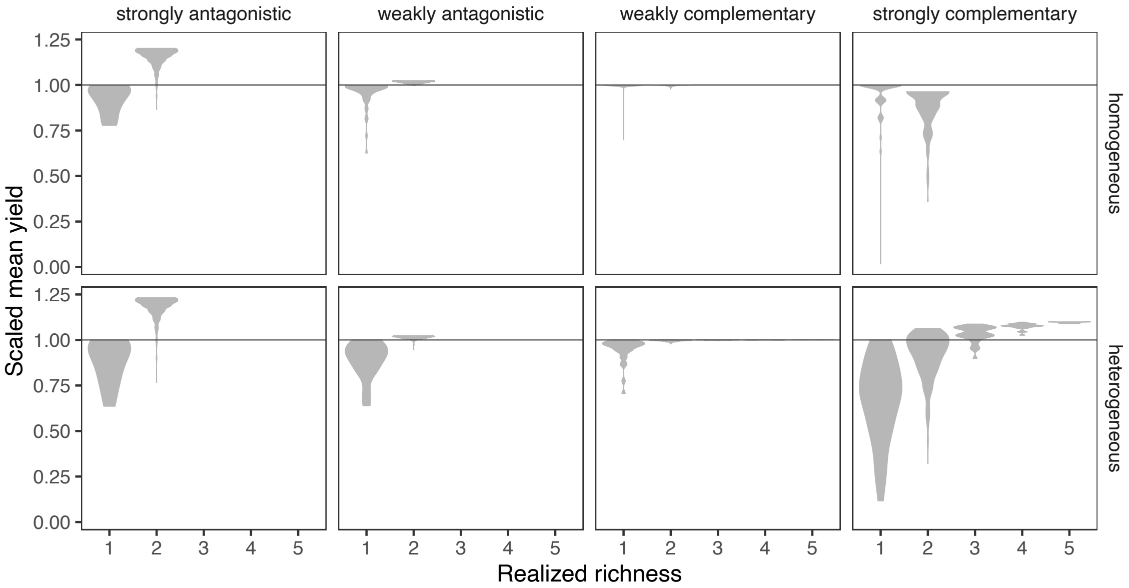

**Figure S1**. Relationship between yield and realized richness for the 8 scenarios. The small light points represent the equilibrium community resulting from seeding all possible subsets of the global species pool. The large darker points give the average yield and realized richness, where the average is taken across all possible communities for a given level of introduced richness.

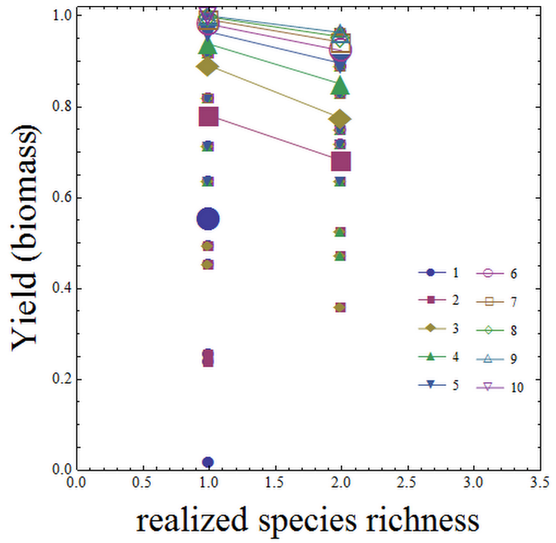

**Figure S2**. Relationship between realized richness and yield for the strongly complementary case in a homogeneous environment (scenario 7, Table 1). Each small point represents an equilibrium community resulting from seeding a given subset of the global species pool, coded by levels of introduced richness (from 1 to 10). For each level of introduced richness, the large points give the average yield across each level of realized richness. The lines connecting these points highlight the fact that within scenario 7, total biomass is negatively related to realized species richness for any given level of introduced richness.

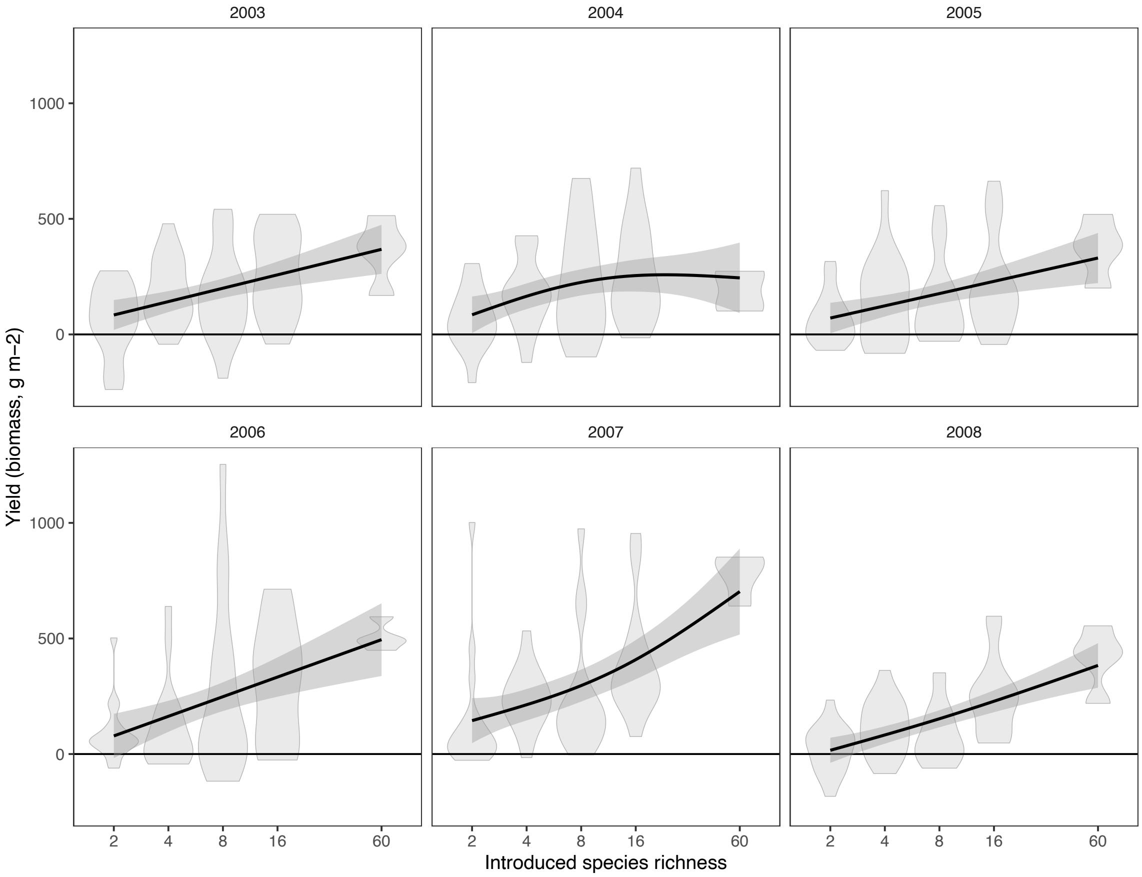

**Figure S3**. Jena yield (biomass) over time, relative to hypothetical community based on monoculture performance, as a function of introduced richness (log scale). Complementarity-Selection analysis (Fig. 6) partitions these responses.

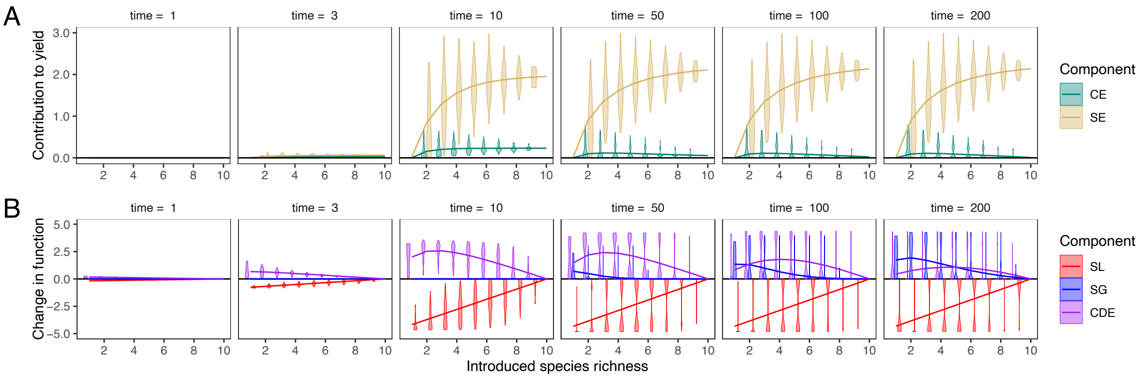

**Figure S4**. Transient dynamics in Complementarity-Selection effects and Community Assembly components as assembly proceeds towards equilibrium in the scenario where species compete for heterogeneous, strongly complementary resources.

Appendix A: **Modeling the different resource scenarios**

In this Appendix, we present how the different resource scenarios are modeled**.** This is done used a flexible form of the $g_j(R_1, R_2)$ function developed by Schreiber and Tobiason (2003):

$$g_j(R_1, R_2) = [(x_{1j}R_1)^\alpha + (x_{2j}R_2)^\alpha]^{\frac{1}{\alpha}} \quad \text{(1A)}$$

Here $\alpha$ is a tunable shape parameter that determines the type of resource (i.e. complementary or antagonistic). The traits $x_{1j}$ and $x_{2j}$ are species *j*'s acquisition rates for $R_1$ and $R_2$, and control how efficient species *j* is at acquiring resources. We modeled the four different resource scenarios by picking four different values for the shape parameter $\alpha$ : strongly complementary ($\alpha \ll 1$), strongly antagonistic ($\alpha \gg 1$), weakly complementary ($\alpha \lesssim 1$) and weakly antagonistic ($\alpha \gtrsim 1$). To fully specify the model, we also had to tune species' consumptive impacts on their resources, $\boldsymbol{C_{ij}(R_1, R_2)}$. To do so we followed Koffel et al. (2016) and modeled per capita consumptive impacts separately for each resource scenario to better match the biology of resource uptake, as described below.

For strongly complementary resources ($\alpha \ll 1$), we assumed that both resources are necessary to grow, which means that growth translates into proportional uptakes of both resources through fixed stoichiometric coefficients $q_{ij}$:

$$C_{ij}(R_1, R_2) = q_{ij}\, g_j(R_1, R_2) \quad \text{(2A)}$$

This is consistent with the way essential resources are usually modeled (Tilman 1982). We further assume that $q_{1j} = x_{2j}$ and $q_{2j} = x_{1j}$, which ensures that every species has a greater impact on the resource that most limits its growth, a standard necessary condition for coexistence to happen in resource competition theory (Tilman 1982, Leibold 1995, Chase and Leibold 2003).

For strongly antagonistic resources ($\alpha \gg 1$), resources can be substituted for each other, but a mixed uptake does not translate additively into growth because of the antagonism between the two resources. Such antagonism can for example happen when the two resources are patchily distributed. In this scenario, the consumers focus their uptake on the most profitable resource (based on its abundance) and avoid the least profitable (often rarest) one. Mixed uptake only occurs for a narrow range of availabilities corresponding to equally profitable resources, where the consumer is about to switch from consuming one resource to consuming the other. Switching uptake can be modeled by the following mathematical expression:

$$C_{ij}(R_1, R_2) = \frac{(x_{ij}R_i)^\alpha}{(x_{1j}R_1)^\alpha + (x_{2j}R_2)^\alpha}\, g_j(R_1, R_2) \qquad \text{(3A)}$$

This choice of consumptive impact is consistent with mass balance of substitutable resources, where all the resources taken up translate into growth by an equivalent amount of biomass. This can formally be checked by summing the two uptake rates of a species $j$ given by Equation (3A) and retrieving its growth rate ($g_j = C_{1j} + C_{2j}$).

Finally, in the weakly complementary and weakly antagonistic resource scenarios ($\alpha \lesssim 1$ and $\alpha \gtrsim 1$, respectively), resources are close to being consumed proportionally to their availability, i.e $C_{ij}(R_1, R_2) \approx x_{ij}R_i$, consistent with their close proximity to the perfectly substitutable case ($\alpha = 1$; Schreiber and Tobiason 2003). Such a close-to-proportional consumptive impact is obtained by borrowing equation (3A) from the antagonistic resource scenario and using it for these two weakly interacting scenarios. Indeed, equation (3A) tends towards linear consumption when $\alpha$ tends towards 1.

To generate a species pool with varying competitive abilities on the two resources, we also implemented a trade-off between the traits $x_{1j}$ and $x_{2j}$ of each species, which control how efficient species $j$ is at acquiring each resource. We assumed these two traits to be bound by a linear trade-off for simplicity ($x_{1j} + x_{2j} = 1$). Varying these two traits between 0 and 1 generates a generalist-specialist continuum.

Appendix B: **Facilitation model**

To study the effects of facilitation on the Complementarity-Selection effects and the Community Assembly components, we developed a variant of the resource competition model from the main text specifically tailored to model the interactions between nitrogen-fixing and non-fixing terrestrial plants. This model is very similar to the model of Koffel et al. (2018), with a few differences that will be highlighted below. This model still fits the general form of equation (1), with now $R_1$ and $R_2$ respectively being soil Phosphorous (P) and Nitrogen (N). Yet, the expressions of both the per capita growth rate and the consumptive impacts differ from the general form from Schreiber and Tobiason (2003) presented in the main text. The growth rate $g_j(R_1, R_2)$ here reads:

$$g_j(R_1, R_2) = \mathrm{Min}(x_{1j}R_1, x_{2j}R_2 + f_j)$$

As the two resources $R_1$ and $R_2$ correspond to P and N, they are modeled as being strictly essential ($\alpha = -\infty$ in the approach presented in the main text) which results in taking the minimum of the growth rates on the two resources via the "Min" function.

The extra term that is not accounted for by the Schreiber and Tobiason (2003) approach is the constant fixation rate $f_j$ that boosts growth when N is limiting. The fixation rate $f_j$ is the focal trait here, and the species pool was generated by selecting species with varying values of $f_j$. A species with $f_j = 0$ is a non-fixer, while species with non-zero values of $f_j$ describe N-fixers with various intrinsic abilities to fix N. We included a cost to N-fixation in the form of a decreasing ability to acquire phosphorous through $x_{1j}$ as $f_j$ increased, mathematically given by $x_{1j} = h(f_j)$, with $h$ a decreasing function. Conversely, soil nitrogen acquisition $x_{2j}$ was assumed to be the same for all species, and thus taken constant.

For facilitation to happen, N fixed by the N-fixers has to become available to the other species. This is done by modifying the consumptive impacts to account for N restitution through recycling, potentially leading to net negative consumptive impacts when N restitution by an N-fixer is greater than N uptake. This leads to the following consumptive impacts on N:

$$C_{2j}(R_1, R_2) = q_{2j}\left[\mathrm{Min}\left(x_{1j}R_1, x_{2j}R_2\right) - \lambda_{2j}m\right]$$

The positive term in $C_{2j}(R_1, R_2)$ accounts for soil N uptake through growth, and is thus equal to $g_j(R_1, R_2)$ without the fixation term $f_j$ because fixed N comes from the atmosphere, not the soil. The negative term is N restitution through recycling from dead plant biomass (generated at the rate $m$, assuming the same mortality rate for all species for simplicity) with an efficiency given by $\lambda_{2j}$. We assumed that the efficiency $\lambda_{2j}$was larger for species with large fixation rate $f_j$, translating the fact that N-fixing species tend to see their dead biomass more readily recycled. The coefficient $q_{2j}$ is a stoichiometric coefficient similar to the ones introduced in the models from the main text.

On the P side, the consumptive impact reads:

$$C_{1j}(R_1, R_2) = q_{1j}[g_j(R_1, R_2) - \lambda_{1j} m]$$

The first term of the sum is here exactly equal to growth $g_j(R_1, R_2)$, which translates the fact that unlike N, all the P entering plant biomass has been taken up from the soil. We assume that P is also recycled, through an efficiency $\lambda_{1j}$ assumed to be the same for all species for simplicity. The coefficient $q_{1j}$ is again a stoichiometric coefficient.

The global species pool was generated by selecting 10 equally spaced values for the fixation rate $f_j$, and the corresponding phosphorous acquisition rates $x_{1j}$ and N-recycling efficiencies $\lambda_{2j}$ given by the trade-offs detailed above. All the other parameters were chosen to be the same for these 10 species.

For every given subset of the species pool, community assembly was simulated numerically by seeding the species present in that subset altogether and letting them grow until an equilibrium was reached.

Because N-fixing species with large $f_j$ can, under certain circumstances, have negative consumptive impacts $C_{2j}(R_1, R_2)$, their presence leads to the net enrichment of the N pool. This results in strong fixers facilitating non-fixers and weak fixers (species that have small fixation rates $f_j$) through two mechanisms: 1) a strong N-fixer can coexist at equilibrium with a weak fixer, supplying the latter with limiting N, preventing the weak fixer from going extinct or enabling it to reach a higher biomass than in the absence of the strong fixer, and 2) in a initially very N-limited environment that prevents the weak-fixer from establishing, a strong N-fixer can establish and lead to soil N accumulation. This makes establishment of the weak fixer possible, but eventually leads the strong fixer to its own demise as the weak fixer excludes it. This second mechanism is a pure effect of community assembly that can generate facilitation-driven succession, as described in detail in Koffel et al. (2018).
